# Assessment of abduction movement in older people with painful shoulder: an analysis based on inertial sensors

**DOI:** 10.1101/398123

**Authors:** Cristina Roldán-Jiménez, Jaime Martín-Martín, Paul Bennett, Steven McPhail, Antonio I Cuesta-Vargas, Graham K. Kerr

## Abstract

Reduced range of motion in the shoulder can be a source of functional limitation. Current quantitative evaluation systems are limited to assessing the functionality or the maximum articular amplitudes in each of the planes of movement, both in isolation. These separate clinical evaluation systems may not allow the identification of the underlying impairments contributing to the functional limitation. The use of inertial sensors to quantify movement in addition to more common clinical assessments of the shoulder may allow clinicians to understand that are potentially unnoticed by the human eye. The main objective of this cross-sectional study was to generate an explanatory model for shoulder abduction based on data from inertial sensors. Shoulder abduction of thirteen older adults suffering from shoulder dysfunction was evaluated using two inertial sensors placed on the humerus and scapula. Movement variables (maximum angular mobility, angular peak of velocity, peak of acceleration) were used to explain the functionality of the upper limb (assessed using the Upper Limb Functional Index). Abduction movement of the shoulder was explained by six variables related to the mobility of the shoulder joint complex. A manova analysis was used to explain the results obtained on the functionality of the upper limb. The MANOVA model based on angular mobility explained 69% of the variance of the ULFI value (r-squared=0.69). The most relevant variables were the abduction-adduction of the humerus and the medial and lateral rotation of the scapula. However, given the limited sample size, none of these individual variables were statistically significant in the decomposition model on their own. The method used in the present study reveals the potential importance of the analysis of the scapular and humeral movements for comprehensive evaluation of the upper limb. Further research should include a wider sample and may seek to use this assessment technique in a range of potential clinical applications.

## Introduction

Shoulder disorders are highly prevalent in the population and to a greater extent among older adults; in many cases, biomechanical anomalies are asymptomatic [1,2]. Despite the potential absence of pain, limitations caused by a reduced range of motion (ROM) can have a detrimental impact on the performance of activities of daily living [3]. Clinically, the evaluation of patients’ daily functioning in their upper limbs is often based on self-administered questionnaires, such as Disability of the Arm Shoulder and Hand (DASH) [4] or Upper Limb Functional Index (ULFI) [5] and functional physical assessments [6]; due to the high cost of the imaging devices such as motion tracking with multiple camera system or real-time ultrasound.

In this regard, there is increasing evidence supporting the combined use of new technologies with functional physical tests or ROM assessments for more comprehensive biomechanical diagnostics. Technologies used for this purpose include: X-ray [7–9], magnetic resonance imaging [10], 3D imaging models [11] and inertial sensors [2,12], among others. Currently, one of the devices most frequently used for clinical purposes is inertial sensors due to their small size, reliability, and accuracy for registering human movement (speed, acceleration, and orientation) [13].

For the assessment of shoulder mobility with inertial sensors, it is necessary to use two or three inertial sensors located on the skin adjacent to the humerus, scapula, and chest [14]. Inertial sensors have been used alongside other objective assessments of the quality and quantity of movement in patients with chronic painful shoulders [15,16]. Similarly, these sensors have sufficient sensitivity to discriminate between healthy and affected subjects, complementing the results of standardized assessment scales [15–19].

Changes in shoulder mobility, whether symptomatic or asymptomatic, may be due to bone and muscle-related disorders affecting the rotator cuff [20,10,9,11,8]. This may limit upper limb functionality and, in some cases, is directly associated with age [20]. Prior research has indicated there are no significant differences in the neutral positioning of the shoulder in symptomatic and asymptomatic subjects. However, significant changes in internal mobility of the shoulder joint complex have been detected when the shoulder flexes above 90 degrees [18]. This is because the scapula has movement in three spatial planes [21], which can change the orientation of the arm[7,9,11,19]. In addition to the scapula, shoulder muscles also change their activation level according to the speed and ROM of flexion or abduction[22].

Positive correlations between acceleration values measured with inertial sensors and function-related questionnaire responses (DASH) have been reported previously. Higher acceleration values reflected greater shoulder functionality [16]. The asymmetry of shoulder movement assessed with inertial sensors has also been correlated with patient-reported functionality; greater asymmetry indicated less functionality [23].

Therefore, describing an approach for assessing shoulder mobility based on inertial sensors in subjects who have functional limitations is an important next step in advancing the field. Ideally, the assessment with multiple inertial sensors will discriminate whether the displacement of body segments is below or above the normal values and contribute to functional limitation; providing additional information for clinical use. Therefore, inertial sensors could be used to compliment more traditional assessments when diagnosing shoulder dysfunction. The main objective of this study was to design a multivariate model for upper limb dysfunction based on inertial sensors, thereby obtaining predictors of upper limb dysfunction based on shoulder movements.

## Materials and methods

### Subjects

A cross-sectional study was designed to evaluate abduction movement of the shoulder with two IntertiaCube3 Sensors [24]. Thirteen participants (9 females, 4 males) were recruited from a specialized orthopedics clinic. They had previously been diagnosed with rotator cuff tears by magnetic resonance imaging and were on the waiting list for surgical intervention. Inclusion criteria were age between 18 and 75 years old, Body Mass Index (BMI) between 18 and 42 and presence of a confirmed rotator cuff tear. Participants were excluded if they declined to participate in the study or had concurrent or alternative etiologies for their shoulder dysfunction.

An a-priori sample size of 9 participants was calculated for an α error of 0.05, a statistical power of 0.8 and β error of 0.7, based on data from a systematic review on the use of inertial sensors to measure human movement [25].

The study was approved by the ethics committee of the University of Malaga (Faculty of Health Sciences) and complied with the principles of the Declaration of Helsinki [28]. All participants provided informed consent prior to taking part in the study.

### Apparatus

Two IntertiaCube3 Sensors (Billerica,MA,US) [24] were used to measure shoulder abduction of each subject. This sensor has high performance: 4ms latency, 180 Hz rate, high accuracy (below 1° of error), 3 degrees of freedom (Yaw, Pitch, and Roll) and a little size and weight. Acceleration (m/s^2^) and angular mobility (°) of shoulder abduction were measured with these sensors in the three spatial axes (Z-Yaw, Y – Pitch, X - Roll); each spatial axis was related to the movement of the corporal segment (Table 1).

**Table 1.** Real movement by axis and plane.

| Region | Axis- plane | Real Movement |
| --- | --- | --- |
| **Humerus** | X -Roll | FL–EX |
|  | Y - Pitch | IN–EX |
|  | Z - Yaw | AB–AD |
| **Scapula** | X - Roll | AN–PO |
|  | Y - Pitch | PR–RE |
|  | Z - Yaw | ME–LA |

Two inertial sensors were placed on the humerus and the scapula following the protocol designed by Cutti et al. (2008) [14]. To ensure the correct measurement data, the skin of participants was cleaned with alcohol before attaching the sensors to the skin with double-sided adhesive.

Before making the recordings, inertial sensors were calibrated to 0 following the protocol established by the manufacturer’s software [26]. This software was the same as used for recording data; a low-pass filter (Kalman filter) was applied while recording data.

### Procedure

Prior to registration made with inertial sensors, patient characteristics including the Spanish version of Upper Limb Functional Index (ULFI) [5]. ULFI is an upper extremity outcome measure that consists of a 25-item scale that can be transferred to a 100-point scale. It also has strong psychometric properties [27]. Its Spanish version has demonstrated high internal consistency (α = 0.94) and reliability (r = 0.93) [5]. Placed in the standing position and with the upper extremity in the neutral position, participants performed three full shoulder abductions. That is, lifting the arm sideways until the hand reaches as high as possible. Two sets of three repetitions were recorded; the second repetition of each set was the one chosen to be analysed. Patients were told to perform shoulder abduction at a natural speed until their movement had reached its end of range of motion. Questionnaires were recorded. Abduction of the affected shoulder was measured on the humeral and scapular sections. The main variables analyzed were: maximum angular mobility (°), angular peak of velocity (°/s), peak of acceleration (m/s^2^).

### Data analysis

Descriptive analyses (mean, SD) were used for participant characteristic variables (weight, height, BMI, and age) to describe the sample. Based on the yaw, pitch and roll values obtained by the sensor the minimum peaks obtained by each sensor were subtracted from the maximum peaks for each of the variables (acceleration and speed); the norm of the resultant vector (Nrv) was calculated 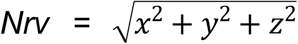 in order to obtain the mean speed and acceleration of the movement performed. The means of peak of maximum angular mobility inside the abduction movement in each of the 3 axes (yaw, pitch and roll) were used in order to create an ANOVA multivariate analysis (MANOVA) model.explaining the ULFI questionnaire results.

## Results

Functional impairment of the participants was reflected by high ULFI values (mean±SD) 70.96±20.93 [5]. The anthropometric characteristics of the participants were age 52.68±9.78 years, weight 75.58±17.98 kg, height 1.64±0.09 m and body mass index 28.22±6.59 kg/m^2^.

The mean maximum angular mobility in humerus abduction axis (72.97°) indicated that a substantial reduction in shoulder mobility was present in the sample. The reductions of the range movement, also affect to scapular section, where the mean range was 12.63° for protraction-retraction (Table 2).

**Table 2:** Mean (95%*CI*) sensor peak maximum of angular mobility (°) from each axis in abduction movement.

| Surface placement | HUMERUS |  | SCAPULA |  |
| --- | --- | --- | --- | --- |
|  | Motion | Mean (95%CI) | Motion | Mean (95%CI) |
| **X** | IN-EX | 38.0 (16.17-59.85) | AN-PO | 15.60 (2.33-13.86) |
| **Y** | AB-AD | 77.56 (48.18-106.92) | PR-RE | 12.63 (5.56-19.70) |
| **Z** | FL-EX | 35.72 (21.12-50.31) | ME-LA | 4.57 (1.87-7.28) |

. The MANOVA model explained 69% of the variance of the ULFI value. In an exploratory decomposition of the multivariate model, the main explanatory variable was the value of the humerus AB-AD movement (*p*=0.093) followed of the scapular ME-LA (*p*=0.195). However, given the limited sample size in the present study, none of these individual variables were statistically significant in the decomposition model on their own. On the other hand, there was less (or no) indication that gender and scapular anterior–posterior tiling movement were likely to have importance for the explanation of the model (Table 3).

**Table 3:**
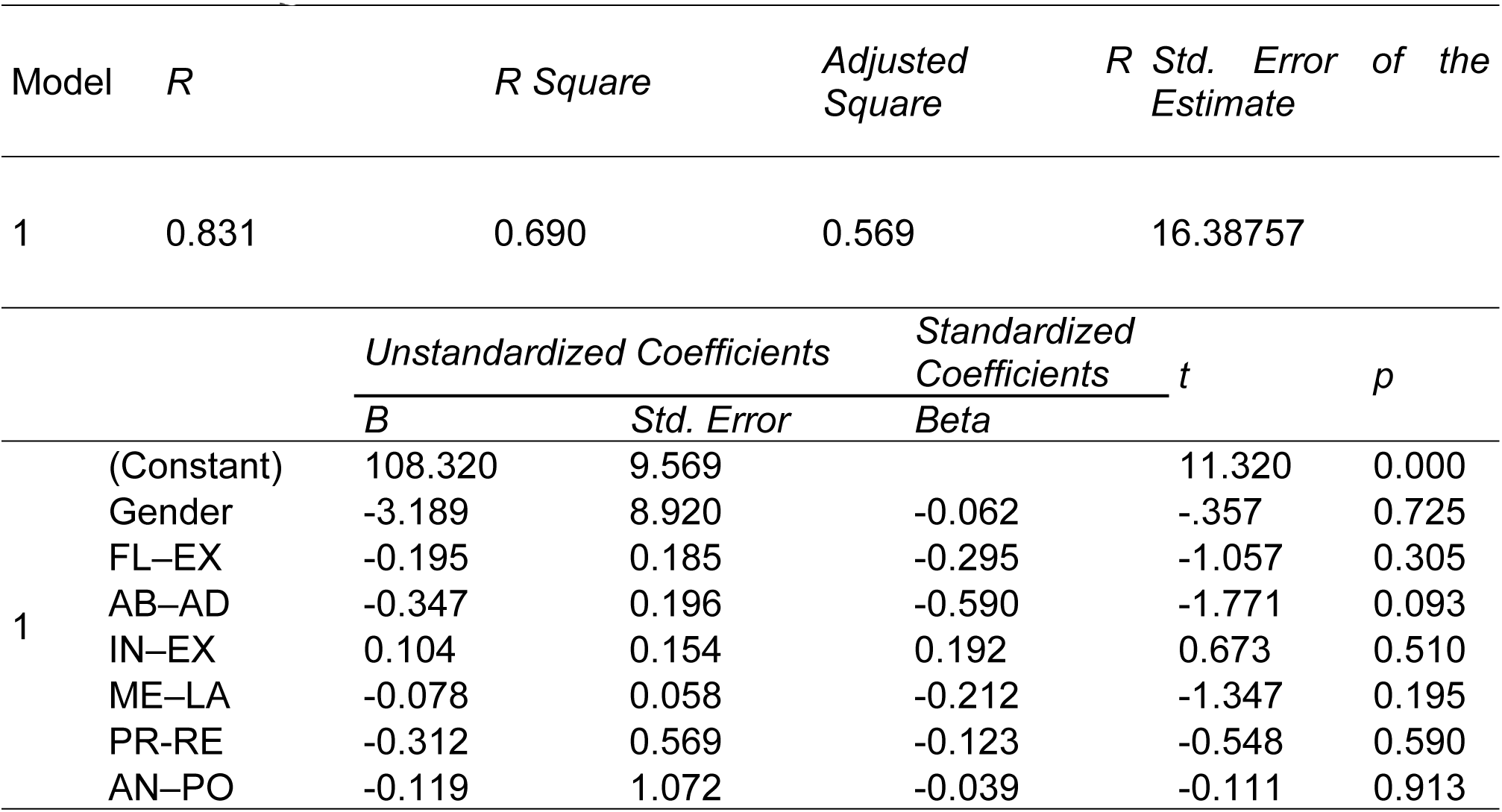
Decomposition of the multivariate model.

The mean (95%CI) peak of acceleration (m/s^2^) and velocity (°/s) and peak from norm of the resultant vector in abduction is presented in Table 4.

**Table 4.**
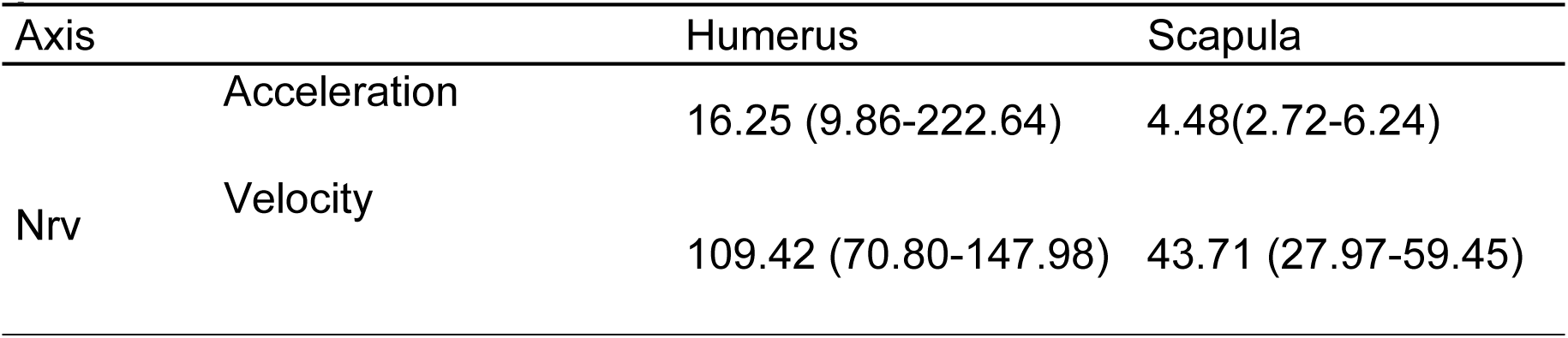
Mean (95%CI) peak of acceleration (m/s^2^) and velocity (°/s) and peak from norm of the resultant vector in ABD.

## Discussion

Shoulder abduction has been described by the use of two inertial sensors in the present study. In this case, they were placed on the scapula and humerus. The results obtained were used in order to create a multivariate model for the description of abduction movement(s) that explained shoulder function as reported by patients using the ULFI questionnaire. Thus, the overall objective of the study was fulfilled. The maximum values of acceleration and angular velocity refer to the normal movement performed by the participants (they were asked to perform shoulder abduction at normal speed). This requirement allowed us to create a model that was more faithful to shoulder movement in daily living. In accordance with the results obtained, the inertial sensors have potential for complementary description and quantification of perceived disability in the upper limb; because of the results of MANOVA model (r-squared=0.69).

Prior descriptions of shoulder abduction have been carried out by various authors in recent decades [7–11,20]. Most of these prior studies focused on rotator-cuff fatigue although some elements of biomechanical mobility during shoulder abduction have been examined using X-rays [7–9], ultrasound techniques [20], magnetic resonance [10] or computed tomography [11]. Unfortunately these previously reported approaches to examining biomechanical shoulder abduction movement are likely to be too costly for daily use in clinical settings and the inability to follow the movement in real time. However inertial sensors do not have these limitations [13].

The inertial sensors used in the present study allowed an assessment based on angles, speed and acceleration movements. However, they do not provide the same level of insight pertaining to osseous structures involved in the movement as the aforementioned imaging technologies. Nonetheless, the supplementary biomechanical information provided by inertial sensors may add value in the context of biomechanical diagnostics in clinical settings.

According to findings from the movement decomposition model reported in Table 3, humerus ABD-ADD and scapula AN-PO may be the movement components that have the greatest association with self-reported upper limb function. This finding was not surprising; however, this was accompanied in greater measure for the scapular anterior-posterior movement rather than lateralization (that is to say, scapular ME-LA). These findings from the present study are consistent with a previous study that focused on the scapulothoracic joint which found a reduced ME-LA motion during elevation in symptomatic subjects [29]. In contrast with these results, another previous study that examined 3D scapula kinematics using Polhemus Fastrak found an increased scapular lateral rotation as a compensatory pattern in pathological shoulders [30]. A potential explanation for this observation is that the middle deltoid is the muscle that performs arm elevation with greater activity over 75° of shoulder abduction, while the supraspinatus is more effective at low angles [11]. Likewise, Duc et al. (2014) observed different levels of muscle activation for shoulder abduction in the same subject depending on the affected or healthy side [31].

In this regard, the rotator cuff has an important role in shoulder abduction[9]. Prior studies have shown tears in the rotator may or may not be symptomatic or associated with functional deficits, and they are positively associated with older age [10]. Several studies have demonstrated that shoulder mobility does not maintain a direct relationship with the size or thickness of the tear [10,20]; and in some cases there may be accommodation of the humeral head in the glenoid [8,9,11].

In the present study, the findings presented in the Table 3 ANOVA model may represent something of the accommodation or biomechanical adaptation of the scapula and humerus movement using the two additional planes of movement rather than abduction alone. This is consistent with prior reports from other authors regarding the 3-dimensional movement of the scapula during performance of shoulder abduction in symptomatic and asymptomatic subjects [21]. Although the 3 dimensional movement of the scapula has been discussed in prior research, the symptomatic nature of participants in the present study (mean ULFI 70.96) and quantification of 3-dimensional movement using a straight-forward inertial sensor setup means that finding from the present study are likely to have particular relevance for this clinical population.

The association between the dysfunctionality of the upper limb (high values in ULFI) and movement of the arm was consistent with other studies. Jolles et al (2011) identified a positive coefficient of correlation (R>0.61) for 3 kinematic variables in 4 different questionnaires functionality. One of the questionnaires tested, the Simple Shoulder Test, obtained an excellent linear correlation (R=0.80) with shoulder power [16]. The results obtained by Körver et al (2014) also showed positive relationships between kinematics asymmetry scores, understood as relative difference between healthy and affected side, and DASH and Simple Shoulder Test Questionnaires (R=0.79); which indicated a high diagnostic power to differentiate between healthy and affected side. These results are consistent with those obtained in the present study in which the values of humeral AB-AD explained almost 60% on the negative direction of the variance of the questionnaire functionality (Table 3). Therefore, a lower level of AB-AD was associated with a higher score obtained on the ULFI (which corresponds to a less functionality).

In clinical contexts, shoulder assessment is usually done by traditional clinical tests that are based on the premise that it is possible to isolate individual structures by compressing or stretching the tissue of interest. However, this is not possible without affecting the state of adjacent structures [32] because rotator cuff tendons are interwoven as a functional unit [33]. For example, some clinical tests that are intended to implicate supraspinatus pathology have been demonstrated (using electromyography) to activate eight or nine other muscles [34]. Hence, the employment of these tests leads to inaccuracy in their findings [32–35].

Furthermore, the clinical expression of shoulder injuries is highly variable [36–38]. Besides clinical test, in shoulder assessment image tests are also employed. However, these too may be considered invalid at times, as there are a large number of asymptomatic individuals who have structural shoulder abnormalities[32]. Hence, at present, the evaluation and diagnosis of joint pathology of the shoulder joint is a complex clinical endeavor prone to uncertainty [33]. Results from the present study reinforce the potential use of both inertial sensor and questionnaires to assess shoulder function building on prior research in the field that has established each of these as independently validated measurement instruments. Research contributions that identify potential predictors of upper limbs dysfunction based on validated instruments may not only have a role in diagnostics, but also have additional potential in quantify the effect of treatment. By extension, this may also lead to further developments that assist in improving predictions of which patients are likely to receive the greatest benefit from surgical or conservative interventions. To build on findings from the present study, future research may also seek to measure the same movement in a person twice (one on each side) and also for patients to report the functionality of each of their upper limbs, as well as their global upper limb functions. A study of this nature would have the potential to correlate the functional capacity of each of the sides with the same kinematics values, while also considering the roles of unilateral or bilateral kinematic deficits and hand dominance on self-reported upper limb functioning to investigate the potential mediating role of the unaffected arm.

## Conclusions

The functionality of the shoulder is a key element in activities of daily living. The application of complementary assessment technologies like inertial sensors to traditional clinical upper limb assessments may add value for the functional diagnosis of patients with shoulder pain. The method used in the present study reveals the potential importance of the analysis of the scapular and humeral movements for comprehensive evaluation of the upper limb. The individual analysis of the planes of movement demonstrated the importance of considering the relative contribution of each joint movement. Use of wireless 3-dimentional sensors permitted the consideration of shoulder abduction as a combination of movements dependent on each other within the joint complex of the shoulder. Further research may seek to use this assessment technique in a range of potential clinical applications.

## Acknowledgments

The authors are grateful to the volunteers for their participation and to Mariano Jaimez for his help in data analysis. SMM is supported by a National Health and Medical Research Council (of Australia) fellowship. This study received a grant for Best Protocol Research 2013 from the Chartered of Physiotherapy of Andalusia, Spain [Grant for Best Protocol Research 2013, 5369/16P/SG].

## References

1. Gill TK, Shanahan EM, Allison D, Alcorn D, Hill CL. Prevalence of abnormalities on shoulder MRI in symptomatic and asymptomatic older adults. Int J Rheum Dis. 2014;17: 863–871. DOI:10.1111/1756-185X.12476

2. Louwerens JKG, Sierevelt IN, van Hove RP, van den Bekerom MPJ, van Noort A. Prevalence of calcific deposits within the rotator cuff tendons in adults with and without subacromial pain syndrome: clinical and radiologic analysis of 1219 patients. J Shoulder Elb Surg Am Shoulder Elb Surg Al. 2015;24: 1588–1593. DOI:10.1016/j.jse.2015.02.024

3. Burner T, Abbott D, Huber K, Stout M, Fleming R, Wessel B, et al. Shoulder symptoms and function in geriatric patients. J Geriatr Phys Ther 2001. 2014;37: 154–158. DOI:10.1519/JPT.0b013e3182abe7d6

4. Hudak PL, Amadio PC, Bombardier C. Development of an upper extremity outcome measure: the DASH (disabilities of the arm, shoulder and hand) [corrected]. The Upper Extremity Collaborative Group (UECG). Am J Ind Med. 1996;29: 602–608. DOI:10.1002/(SICI)1097-0274(199606)29:6<602::AID-AJIM4>3.0.CO;2-L

5. Cuesta-Vargas AI, Gabel PC. Cross-cultural adaptation, reliability and validity of the Spanish version of the upper limb functional index. Health Qual Life Outcomes. 2013;11: 126. DOI:10.1186/1477-7525-11-126

6. Roe Y, Soberg HL, Bautz-Holter E, Ostensjo S. A systematic review of measures of shoulder pain and functioning using the International classification of functioning, disability and health (ICF). BMC Musculoskelet Disord. 2013;14: 73. DOI:10.1186/1471-2474-14-73

7. Poppen NK, Walker PS. Normal and abnormal motion of the shoulder. J Bone Joint Surg Am. 1976;58: 195–201.

8. Teyhen DS, Miller JM, Middag TR, Kane EJ. Rotator cuff fatigue and glenohumeral kinematics in participants without shoulder dysfunction. J Athl Train. 2008;43: 352–358. DOI:10.4085/1062-6050-43.4.352

9. Yamaguchi K, Sher JS, Andersen WK, Garretson R, Uribe JW, Hechtman K, et al. Glenohumeral motion in patients with rotator cuff tears: a comparison of asymptomatic and symptomatic shoulders. J Shoulder Elb Surg Am Shoulder Elb Surg Al. 2000;9: 6–11.

10. Sher JS, Uribe JW, Posada A, Murphy BJ, Zlatkin MB. Abnormal findings on magnetic resonance images of asymptomatic shoulders. J Bone Joint Surg Am. 1995;77: 10–15.

11. Terrier A, Reist A, Vogel A, Farron A. Effect of supraspinatus deficiency on humerus translation and glenohumeral contact force during abduction. Clin Biomech. 2007;22: 645–651. DOI:10.1016/j.clinbiomech.2007.01.015

12. Louwerens JKG, Sierevelt IN, van Hove RP, van den Bekerom MPJ, van Noort A. Prevalence of calcific deposits within the rotator cuff tendons in adults with and without subacromial pain syndrome: clinical and radiologic analysis of 1219 patients. J Shoulder Elb Surg Am Shoulder Elb Surg Al. 2015;24: 1588–1593. DOI:10.1016/j.jse.2015.02.024

13. Cuesta-Vargas A, Galán-Mercant A, Williams J. The use of inertial sensors system for human motion analysis. Phys Ther Rev. 2010;15: 462–473. DOI:10.1179/1743288X11Y.0000000006

14. Cutti AG, Giovanardi A, Rocchi L, Davalli A, Sacchetti R. Ambulatory measurement of shoulder and elbow kinematics through inertial and magnetic sensors. Med Biol Eng Comput. 2008;46: 169–178. DOI:10.1007/s11517-007-0296-5

15. Duc C, Farron A, Pichonnaz C, Jolles BM, Bassin J-P, Aminian K. Distribution of arm velocity and frequency of arm usage during daily activity: objective outcome evaluation after shoulder surgery. Gait Posture. 2013;38: 247–252. DOI:10.1016/j.gaitpost.2012.11.021

16. Jolles BM, Duc C, Coley B, Aminian K, Pichonnaz C, Bassin J-P, et al. Objective evaluation of shoulder function using body-fixed sensors: a new way to detect early treatment failures? J Shoulder Elb Surg Am Shoulder Elb Surg Al. 2011;20: 1074–1081. DOI:10.1016/j.jse.2011.05.026

17. Lawrence RL, Braman JP, Laprade RF, Ludewig PM. Comparison of 3-dimensional shoulder complex kinematics in individuals with and without shoulder pain, part 1: sternoclavicular, acromioclavicular, and scapulothoracic joints. J Orthop Sports Phys Ther. 2014;44: 636–645, A1-8. DOI:10.2519/jospt.2014.5339

18. Lawrence RL, Braman JP, Staker JL, Laprade RF, Ludewig PM. Comparison of 3-dimensional shoulder complex kinematics in individuals with and without shoulder pain, part 2: glenohumeral joint. J Orthop Sports Phys Ther. 2014;44: 646–655, B1-3. DOI:10.2519/jospt.2014.5556

19. Roren A, Lefevre-Colau M-M, Roby-Brami A, Revel M, Fermanian J, Gautheron V, et al. Modified 3D scapular kinematic patterns for activities of daily living in painful shoulders with restricted mobility: a comparison with contralateral unaffected shoulders. J Biomech. 2012;45: 1305–1311. DOI:10.1016/j.jbiomech.2012.01.027

20. Milgrom C, Schaffler M, Gilbert S, van Holsbeeck M. Rotator-cuff changes in asymptomatic adults. The effect of age, hand dominance and gender. J Bone Joint Surg Br. 1995;77: 296–298.

21. van den Noort JC, Wiertsema SH, Hekman KMC, Schönhuth CP, Dekker J, Harlaar J. Reliability and precision of 3D wireless measurement of scapular kinematics. Med Biol Eng Comput. 2014;52: 921–931. DOI:10.1007/s11517-014-1186-2

22. Castillo-Lozano R, Cuesta-Vargas A, Gabel CP. Analysis of arm elevation muscle activity through different movement planes and speeds during in-water and dry-land exercise. J Shoulder Elb Surg Am Shoulder Elb Surg Al. 2014;23: 159–165. DOI:10.1016/j.jse.2013.04.010

23. Körver RJP, Heyligers IC, Samijo SK, Grimm B. Inertia based functional scoring of the shoulder in clinical practice. Physiol Meas. 2014;35: 167–176. DOI:10.1088/0967-3334/35/2/167

24. InterSense | Precision Motion Tracking Solutions | InertiaCube3TM [Internet]. [cited 10 Aug 2016]. Available: http://www.intersense.com/pages/18/11/

25. Cuesta-Vargas A, Galán-Mercant A, Williams J. The use of inertial sensors system for human motion analysis. Phys Ther Rev. 2010;15: 462–473. DOI:10.1179/1743288X11Y.0000000006

26. InterSense, LLC. Product Manual for use with InertiaCube3 and the InertiaCube Processor.

27. Gabel CP, Michener LA, Burkett B, Neller A. The Upper Limb Functional Index: development and determination of reliability, validity, and responsiveness. J Hand Ther Off J Am Soc Hand Ther. 2006;19: 328–348; quiz 349. DOI:10.1197/j.jht.2006.04.001

28. WMA Declaration of Helsinki - Ethical Principles for Medical Research Involving Human Subjects [Internet]. 19 Oct 2013 [cited 10 Aug 2016]. Available: http://www.wma.net/es/30publications/10policies/b3/

29. Lawrence RL, Braman JP, Laprade RF, Ludewig PM. Comparison of 3-dimensional shoulder complex kinematics in individuals with and without shoulder pain, part 1: sternoclavicular, acromioclavicular, and scapulothoracic joints. J Orthop Sports Phys Ther. 2014;44: 636–645, A1-8. DOI:10.2519/jospt.2014.5339

30. Roren A, Lefevre-Colau M-M, Poiraudeau S, Fayad F, Pasqui V, Roby-Brami A. A new description of scapulothoracic motion during arm movements in healthy subjects. Man Ther. 2014; DOI:10.1016/j.math.2014.06.006

31. Duc C, Pichonnaz C, Bassin J-P, Farron A, Jolles BM, Aminian K. Evaluation of muscular activity duration in shoulders with rotator cuff tears using inertial sensors and electromyography. Physiol Meas. 2014;35: 2389–2400. DOI:10.1088/0967-3334/35/12/2389

32. Lewis JS. Rotator cuff tendinopathy/subacromial impingement syndrome: is it time for a new method of assessment? Br J Sports Med. 2009;43: 259–264. DOI:10.1136/bjsm.2008.052183

33. Lewis J. Rotator cuff related shoulder pain: Assessment, management and uncertainties. Man Ther. 2016;23: 57–68. DOI:10.1016/j.math.2016.03.009

34. Boettcher CE, Ginn KA, Cathers I. The “empty can” and “full can” tests do not selectively activate supraspinatus. J Sci Med Sport Sports Med Aust. 2009;12: 435–439. DOI:10.1016/j.jsams.2008.09.005

35. Hegedus EJ, Goode A, Campbell S, Morin A, Tamaddoni M, Moorman CT, et al. Physical examination tests of the shoulder: a systematic review with meta-analysis of individual tests. Br J Sports Med. 2008;42: 80–92; discussion 92. DOI:10.1136/bjsm.2007.038406

36. Duckworth DG, Smith KL, Campbell B, Matsen FA. Self-assessment questionnaires document substantial variability in the clinical expression of rotator cuff tears. J Shoulder Elb Surg Am Shoulder Elb Surg Al. 1999;8: 330–333.

37. Matthewson G, Beach CJ, Nelson AA, Woodmass JM, Ono Y, Boorman RS, et al. Partial Thickness Rotator Cuff Tears: Current Concepts. Adv Orthop. 2015;2015: 458786. DOI:10.1155/2015/458786

38. Giai Via A, De Cupis M, Spoliti M, Oliva F. Clinical and biological aspects of rotator cuff tears. Muscles Ligaments Tendons J. 2013;3: 70–79. DOI:10.11138/mltj/2013.3.2.070

